# Syntaxin 3-SPI 2 dependent cross-talk facilitates the division of *Salmonella* containing vacuole (SCV)

**DOI:** 10.1101/2022.08.29.505773

**Authors:** Ritika Chatterjee, Nishi Mehta, Subba Rao Gangi Setty, Dipshikha Chakravortty

## Abstract

Intracellular membrane fusion is mediated by membrane-bridging complexes of soluble N-ethylmaleimide-sensitive factor attachment protein receptors (SNAREs). SNARE proteins are one of the key players in the vesicular transport. Several reports shed light on intracellular bacteria modulating host SNARE machinery to establish infection successfully. The critical SNAREs in macrophages responsible for phagosome maturation are Syntaxin 3 (STX3) and Syntaxin 4 (STX4). *Salmonella* actively modulates its vacuole membrane composition to escape lysosomal fusion. A report showed that *Salmonella* containing vacuole (SCV) harbors recycling endosomal SNARE Syntaxin 12 (STX12). However, the role of host SNAREs in SCV biogenesis and pathogenesis is unclear. Upon knockdown of STX3, we have observed a reduction in bacterial proliferation and is restored upon the overexpression of STX3. Post infected live-cell imaging of cells showed STX3 localises to the SCV membranes and thus might help in fusion of SCV with intracellular vesicles to acquire membrane for its division. We also found this interaction abrogated when we infected with SPI-2 encoded T3SS apparatus mutant (STM Δ*ssaV*) but not with SPI-1 encoded T3SS (STM *ΔinvC*). These observations were also consistent in mice model of *Salmonella* infection. Together, these results shed a light on the effector molecules secreted through SPI-2 encoded by T3SS possibly involved in interaction with host SNARE STX3, which is essential to maintain the division of *Salmonella* in SCV and maintenance the principle single bacterium per vacuole.

**Synopsis:** *Salmonella* Typhimurium infection in murine macrophage leads to upregulation of host Syntaxin 3 both at transcript and protein levels at late stage of infection. Syntaxin 3 cross-talk with *Salmonella* containing vacuoles (SCVs) is essential for establishment of replicative niche in host macrophages. The cross-talk between STX3 and SCVs is Salmonella pathogenicity island 2 (SPI-2) dependent and is consistent in mice model of *Salmonella* Typhimurium infection.

## Introduction

Intracellular pathogens are unique in the sense that they have developed numerous strategies to survive and proliferate in their hosts for prolonged periods, and this is achieved by manipulating certain host intracellular trafficking pathways and their components. A large body of literature suggests that the typhoid-causing bacteria *Salmonella* is one such pathogen that successfully establishes an intracellular niche owing to a myriad of virulence effector molecules that it injects into the host cells through type 3 secretion systems (T3SS) encoded by *Salmonella* pathogenicity islands (SPI)-1 and 2 [1-3].

*Salmonella* is a Gram-negative, facultative anaerobic bacterium that primarily infects epithelial cells and macrophages. While the bacteria are phagocytosed by immune cells such as macrophages, they rely on a set of effector proteins translocated by the T3SS-1 for their internalization into epithelial cells. Following its internalization, the intracellular pathogen must ideally be fated for destruction by various phagocytic signaling/immune surveillance systems within the host cell. However, *Salmonella* escapes death by enclosing itself in a host derived membrane-bound vacuole that undergoes a complex series of maturation events to form a specialized compartment that permits its survival and replication. Physiologically, the sequential acquisition of specific Rab proteins on a maturing phagosome regulates dynamic fusion events with host endocytic vesicles, thereby leading to progressive acidification followed by fusion with lysosomes. Even though *Salmonella*-containing vacuoles (SCVs) significantly acquire early and late endosomal markers [4-7], they seem to deviate from the default endo-lysosomal maturation pathway. *Salmonella* is known to effectively ‘remodel’ the vacuole by modulating phosphoinositide metabolism [8, 9] and restricting Rab recruitment [10, 11] on the surface of SCVs, thereby inhibiting lysosomal fusion. *Salmonella* is a particularly interesting pathogen as it resists killing by phagosome acidification instead uses the acidic pH to assemble SPI-2-encoded T3SS, necessary for vacuolar survival inside the macrophages [12, 13]. One of our previous works shows that *Salmonella* resides as a single bacterium per vacuoles, and this facilitates the survival of the bacteria [14]. We have also shown that *Salmonella* downregulates the overall biogenesis of lysosomes[15]; these studies suggest that *Salmonella* stealthily modulates the host endocytic pathways.

An undeniably crucial role in the intracellular survival of *Salmonella* involves fusion events between the SCV and vesicles of the host endocytic pathway. Membrane fusion play a pivotal role in various cellular events like cell signaling, exocytosis, fertilization, neurotransmitter release and is mediated by ‘SNARE’ (soluble N-ethylmaleimide-sensitive factor attachment protein receptors) proteins [16]. SNAREs are structurally classified, as Q-SNAREs (conserved Gln residue) and R-SNAREs (conserved Arg residue). The specific interaction between Q-SNAREs and its cognate R-SNARE results in the formation of a trans-SNARE complex (Qa, Qb, Qc and R), which is responsible for the fusion of two opposing membranes [17, 18].

The intracellular pathogen thrives either in a membrane-bound compartment or in the cytosol and often modulates host endocytic pathways. Recently there have been quite a few advancements in understanding the role of host endocytic machinery in intracellular bacterial infection. *Chlamydia trachomatis* protein IncA mimics SNARE and thus interacts with host SNARE proteins while remaining inside inclusion bodies[19]. In case of *Brucella melitensis* STX4 plays a crucial role in phagocytosis of the pathogen by macrophages[20]. *Legionella pneumophila* type IV effectors *ylfA* and *ylfB* are SNARE-like proteins that form homo-and heteromeric complexes and enhance the efficiency of vacuole remodeling [21]. In *Legionella*, LseA acts as a SNARE protein and has the potential to regulate or mediate membrane fusion events in Golgi-associated pathways[22]. *E. coli* infection hinders the formation of VAMP8-containing exocytic SNARE complexes and thus releases VAMP8-dependent granules by interfering with SNAP23 phosphorylation [23]. There are also a few reports which suggest the crosstalk between SNARE and *Salmonella* infections. It was predicted by an *in-silico* study that *Salmonella* effectors hijack syntaxins by binding to them. Vacuole interactions with endoplasmic reticulum-derived coat protein complex II vesicles modulate early steps of SCV maturation, promoting SCV rupture and bacterial hyper-replication within the host cytosol[24].

*Salmonella* promotes the association of Rab5 to the phagosomes that possibly activate the SNARE to recruitment alpha-SNAP for subsequent binding with NSF to promote fusion of the SCV with early endosomes and inhibit their fusion with lysosomes[5]. *Salmonella* acquires LAMP1 through a SipC-Syntaxin6-mediated interaction to stabilize their niche in macrophages, suggesting other intracellular pathogens might use similar modalities to recruit LAMP1[25]. SNX18 promotes the formation of SCV from the plasma membrane by providing a scaffold to recruit dynamin-2 and N-WASP, and is dependent on the SH3 domain of SNX18. Overexpression of SNX18 increased bacteria internalization, whereas a decrease was detected in SNX18 knocked down cells as well as in cells overexpressing the phosphoinositide-binding mutant R303Q or the ΔSH3 mutant[26]. Syntaxin 8 is involved in the fusion of SCV with early endosomes, and the interaction of SCV localized Syntaxin 4 with SNAP25 mediated the fusion of SCV with infection-associated macro-pinosomes (IAMs) resulting in the enlargement of the vacuole [27, 28].

One *in-silico* analysis has also shown that *Salmonella* effectors can interact with host Syntaxins such as Syntaxin 3, 4, and 12 and thereby modulate its maturation to escape lysosomal fusions [29]. Therefore, we wish to decipher the possible role of the same in *Salmonella* pathogenesis. Syntaxin 3 (STX3) is a SNARE protein involved in vesicle fusion and exocytosis. It is present exclusively on the plasma membrane and is involved in organellar membrane fusion, synaptic vesicle fusion to the pre-synaptic membrane and long-term synaptic potentiation. Unlike its other syntaxin relatives, STX3 is not currently implicated in mediating intracellular infection. In this study, we have demonstrated that the SCV preferentially acquire STX3 on to their surface. We have also shown that the T3SS encoded by SPI-2 is involved in the interaction with STX 3 and that this interaction plays an indispensable role in the division of the SCV.

## Results

### Loss of STX3 in host cells reduces the proliferation of bacteria inside host cells at a late stage of infection

To begin with, our hypothesis of the possible role of host syntaxins in *Salmonella* containing vacuolar (SCV) biogenesis and its impact on further pathogenesis. We have knockdown host STX3, STX4 and STX12 in murine macrophage cell line RAW264.7 using polyethyleneimine (PEI) for 48 hours. To validate the knockdown, we have performed qRT-PCR using primers specific for each syntaxin. The gene expression was normalized to beta-actin as an internal control. About 70-80% knockdown was achieved **(Fig. S1)**. As already discussed, that STX3, STX4 and STX12 could be potential SNAREs having the bacterial pathogenesis host cells[27]. We wanted to delve into the role of host syntaxins in the murine macrophages further. Therefore, we infected RAW264.7 with *Salmonella* Typhimurium (strain 14028) expressing mCherry at an MOI of 25. The cells were then fixed, stained with LAMP1 (as an established SCV marker) and visualized under a confocal scanning laser microscope at different time points. Confocal microscopy images show upon knockdown of host STX3 **(Fig. S2C)**, STX4 **(Fig. S2D)** and STX12 **(Fig. S2E)**, there were no significant differences observed in number of bacteria/ host cells at an early time point of infection (2 hours post-infection) when compared to mock-treated **(Fig. S2A)** or scrambled shRNA treated **(Fig. S2B)**. The data was further quantified **(Fig. S2F)**. Together, these observations suggested that knocking down of STX3, 4 or 12 did not cause any alteration in the phagocytic ability of murine macrophages.

Confocal microscopy images show upon knockdown of host STX3 **(Fig. S3C)** and STX4 **(Fig. S3D)**, there was a significantly lesser number of bacteria/ host cells at the intermediate time-point of infection (6 hours post-infection) when compared to mock-treated **(Fig. S3A)** or scrambled shRNA treated **(Fig. S3B)**. The same was not observed with STX12 knockdown cells **(Fig. S3E)** The data was further quantified **(Fig. S3F)**. These observations suggested that knocking down STX3 and STX4 affected the proliferation of STM WT inside murine macrophages at the intermediate time point of infections.

At the late time-point of infection, confocal microscopy revealed upon knockdown of host STX3 **(Fig. 1C)** that there was a significantly lesser number of bacteria/ host cells at the late time-point of infection (10 hours post-infection) when compared to mock-treated **(Fig. 1A)** or scrambled shRNA treated **(Fig. 1B)**. The same was not observed with STX4 **(Fig. 1D)** and STX12 knockdowns **(Fig. 1E)** The data was further quantified **(Fig. 1F)**. These observations suggested that knocking down of only STX3 and not STX4, and STX12 significantly affects the proliferation of STM WT inside murine macrophages both at intermediate and late time-point of infections. To further confirm our findings, we have stained the knockdown cells with an anti-STX3 antibody. We observed that only in the cells with low expression of STX3 we could see the phenotype of a reduced or lesser number of bacterial cells per host cells (**Fig. 1G**). We have quantified our extent of knockdown in the RAW264.7 cells upon transfection with shSTX3, and we observed around 70% knockdown efficiency in protein level as well **(Fig. 1I)**.

**Figure 1:**
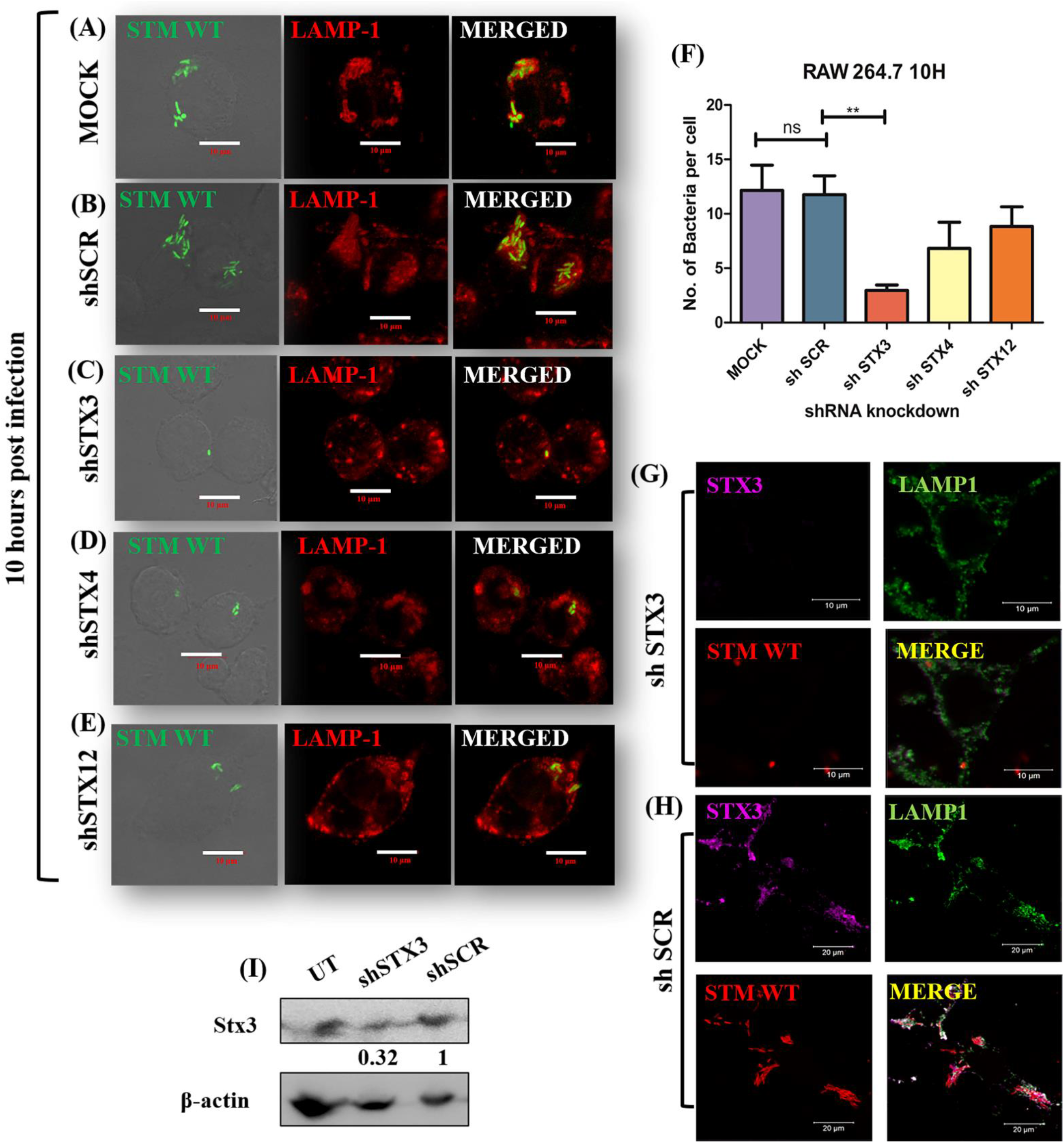
Knockdown of host STX3 leads to reduced no. of bacteria/ host cell at 10 hours post infection with *S*. Typhimurium in murine macrophages RAW 264.7. Representative images of knockdown macrophages infected with *Salmonella* Typhimurium at 10 hours post infection. (A) Mock treated, (B) shSCR (scrambled), (C) shSTX3, (D) shSTX4 (E) shSTX12 and (F) Quantification of number of bacteria/host cell. Confirmation of STX3 knockdown phenotype by immunostaining and immunoblotting; Representative confocal microscopy images of RAW264.7 cells transfected with (G) shSTX3 or (H) shSCR and infected with STM WT and further stained with anti-STX3 antibody; (I) Representative immunoblot of the transfected RAW264.77 cells with shSTX3 or shSCR and compared to untransfected control. (N=3, n= 50 microscopic field) (p<0.05-*, p<0.01-**, p<0.001-***).

### Overexpression of STX3 in murine macrophages (RAW264.7) results in an increased number of bacteria per host cell

To validate the previous observations, we wanted to check the bacterial number/host cell upon infection with STM WT in murine macrophages (RAW264.7) overexpressing rat Syntaxin 3 with a GFP tag (STX3) using confocal imaging. Murine macrophages were transiently transfected for 48h using PEI/FuGENE HD or Lipofectamine 3000 with a plasmid (pEGFP-C1) encoding eGFP-Rat STX3 under the CMV promoter for overexpression. We observed that upon overexpression of STX3 in RAW 264.7 murine macrophages lead to an increase in the number of bacteria/host cells increased significantly as compared to un-transfected at10h post-infection) **(Fig. 2D)**. However, no significant changes were observed at an early time-point (2h post-infection) **(Fig. 2B)**. These observations suggest that upon overexpression of STX3 in murine macrophages as in previous observation, the initial phagocytosis of STM WT is not affected. However, the overexpression of STX3 helps the bacteria to proliferate more inside murine macrophages. The data were quantified as well **(Fig. 2E)**.

**Figure 2:**
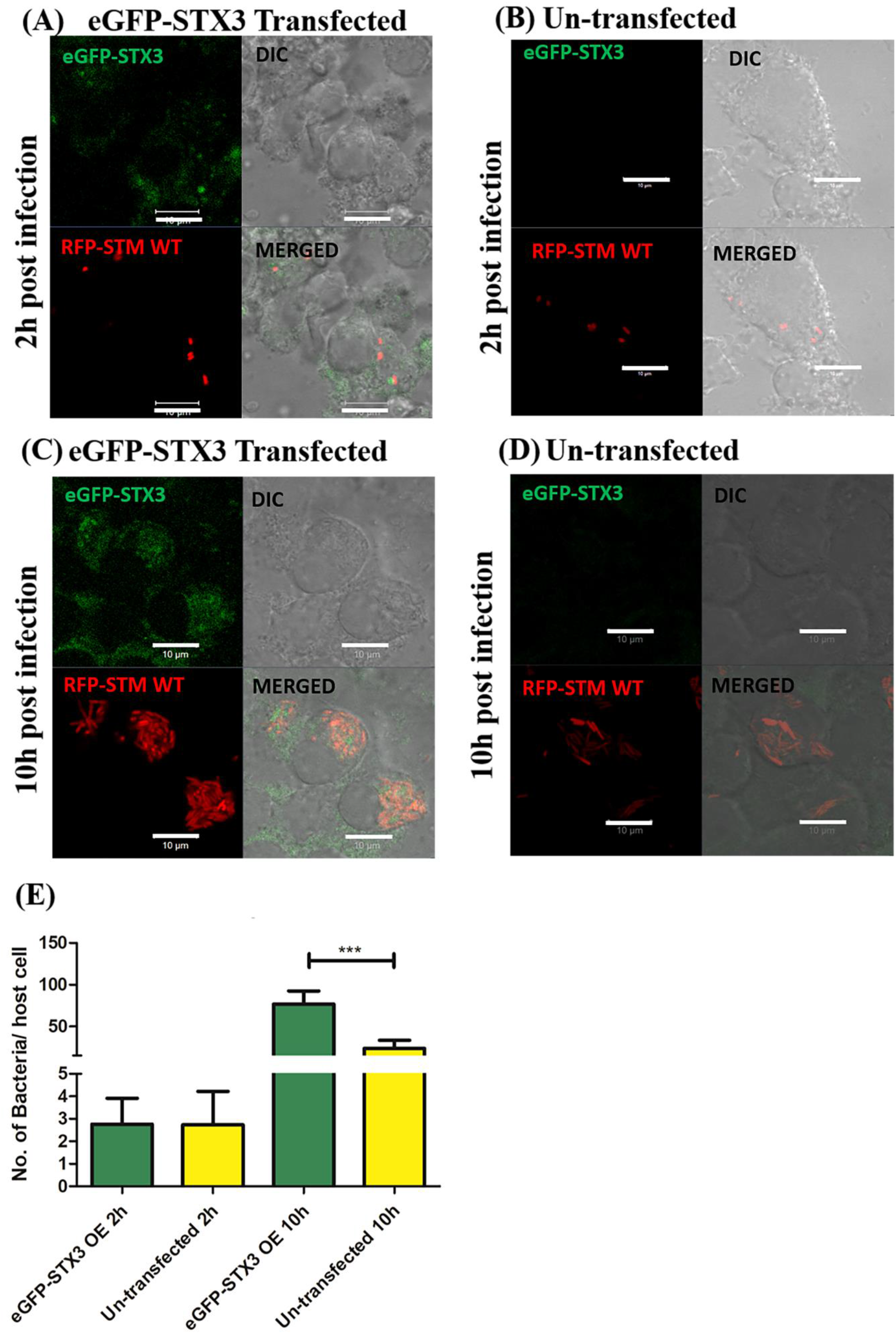
Overexpression of STX3 in murine macrophages (RAW264.7) results in increased number of bacteria/ host cell. Representative images of (A) 2h post infection eGFP-STX3 transfected, (B) 2h post infection un-transfected, (C) 10h post infection eGFP-STX3 transfected, (D) 10h post infection un-transfected, and (E) Quantification of no. of bacterial/host cell (N=2, n= 50 microscopic field) (p<0.05-*, p<0.01-**, p<0.001-***).

### Knockdown of STX3 increases multiple bacteria per SCV in murine macrophages (RAW264.7)

A previous study from our lab has shown that *Salmonella* resides in the host cell as a single bacterium/ vacuole. This gives the pathogen an extra advantage and better survival amidst several host defense mechanisms[14]. It was demonstrated in the same study that the SCV divides along with the bacteria to give rise to two daughter bacterial cells enclosed in individual SCVs[14]. Therefore, it becomes interesting to elucidate the mechanism of SCV division and whether any host protein plays an important role in the same. The SCV enclosing STM has two membranes, one inner membrane of prokaryotic origin of *Salmonella* and one outer phagosomal membrane of eukaryotic origin. Since bacteria have their own machinery to synthesize the cell membrane required during cell elongation and division in two daughter cells. It becomes important for *Salmonella* to hijack or acquire more of the eukaryotic membrane from the endocytic pathway to complete the division of SCV along with the bacteria successfully. SNARE protein plays an important role in membrane fusion in eukaryotic cells. One of our observations suggests that STX3 could be one of the key host proteins that might play a significant role in the acquisition of membrane for SCV division. This is primarily because upon STX3 knockdown, the bacterial number per host cell was greatly reduced compared to mock or scrambled knockdown; few host cells also harbored more no. of bacteria but were sequestered as multiple bacteria/vacuole **(Fig. 3A)**. Also, we have calculated the percentage of host cells having 3 or fewer bacteria and 4 or more bacteria. This analysis has shown a significant difference at 10h time-point were approx. Only 20% of the host cell with STX3 knockdown harbors 4 or more bacteria, among which 80% of them were found to be in multiple bacteria in a vacuole **(Fig. 3A-C)**. To further confirm our finding, we have stained the STX3 knockdown cells with anti-STX3 antibody, and we see similar results **(Fig. S4)**.

**Figure 3:**
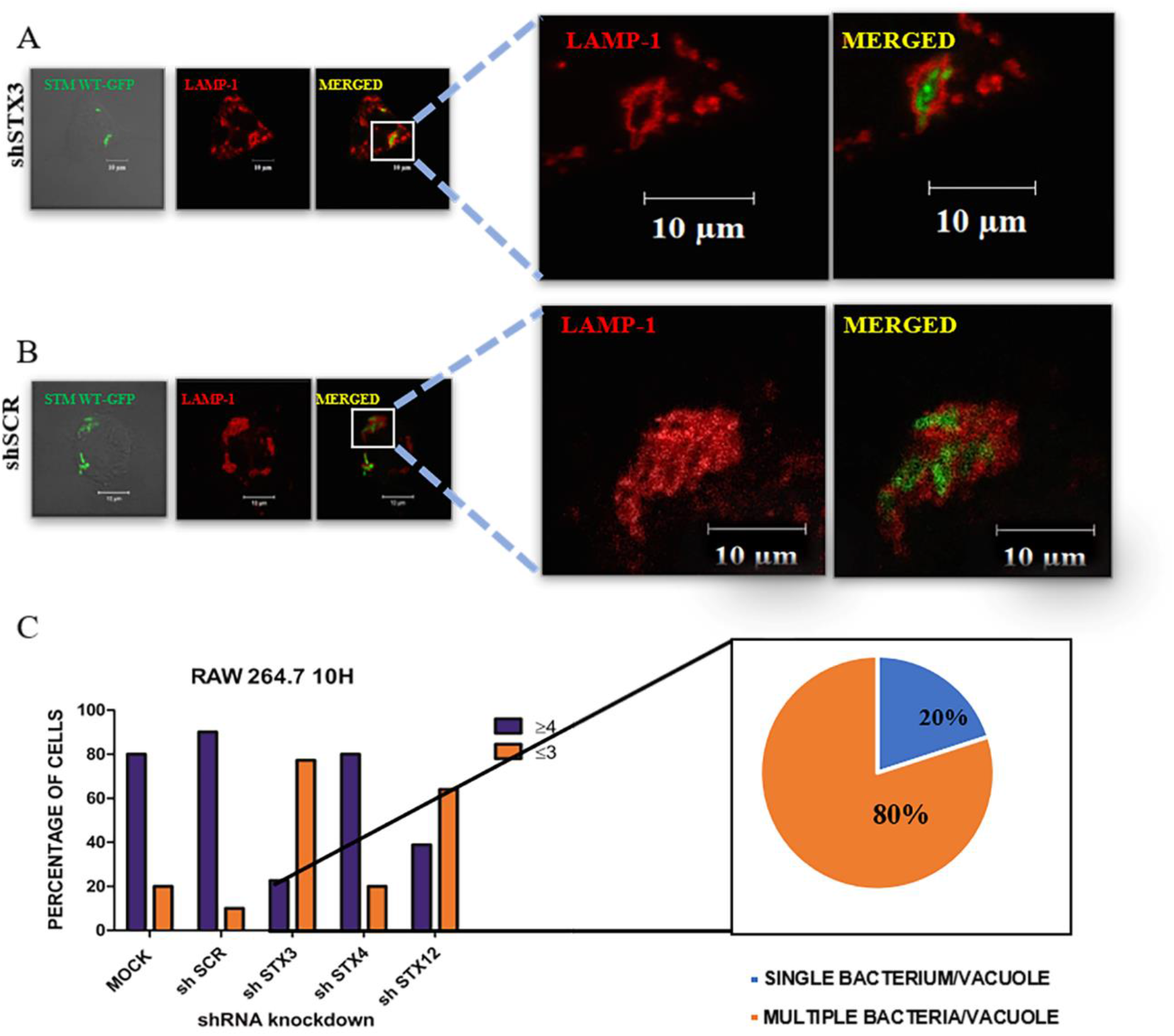
Knockdown of STX3 leads to increase multiple bacteria in a vacuole incidence in murine macrophages (RAW264.7) A) Representative images of multiple bacteria in a vacuole; B) Representative images of scrambled control showing single bacteria in a vacuole; C) Percentage of cell harbouring more than four or three or less bacteria and among them single bacterium per vacuole and multiple bacteria/vacuole percentage. (N=3, n=50 microscopic field).

### At the late time-point of infection with *S*. Typhimurium in murine macrophages (RAW264.7) both transcript and protein level of syntaxin 3 is upregulated

To validate our finding of host STX3 as one of the crucial players in the SCV division, we further wanted to check the levels of STX3 upon STM WT infection at late time points. Murine macrophages were infected with STM WT at an MOI of 10, and cells were harvested for either qRT-PCR or Western blotting to assess transcript and protein levels, respectively. We observed that the levels of host syntaxin 3 are upregulated both at transcript (∼3.8 fold) **(Fig. 4A)** and protein level (∼ 2.5-fold) **(Fig. 4B-C)**, further re-confirming our previous observation of the possible role of STX3 in *Salmonella* pathogenesis.

**Figure 4:**
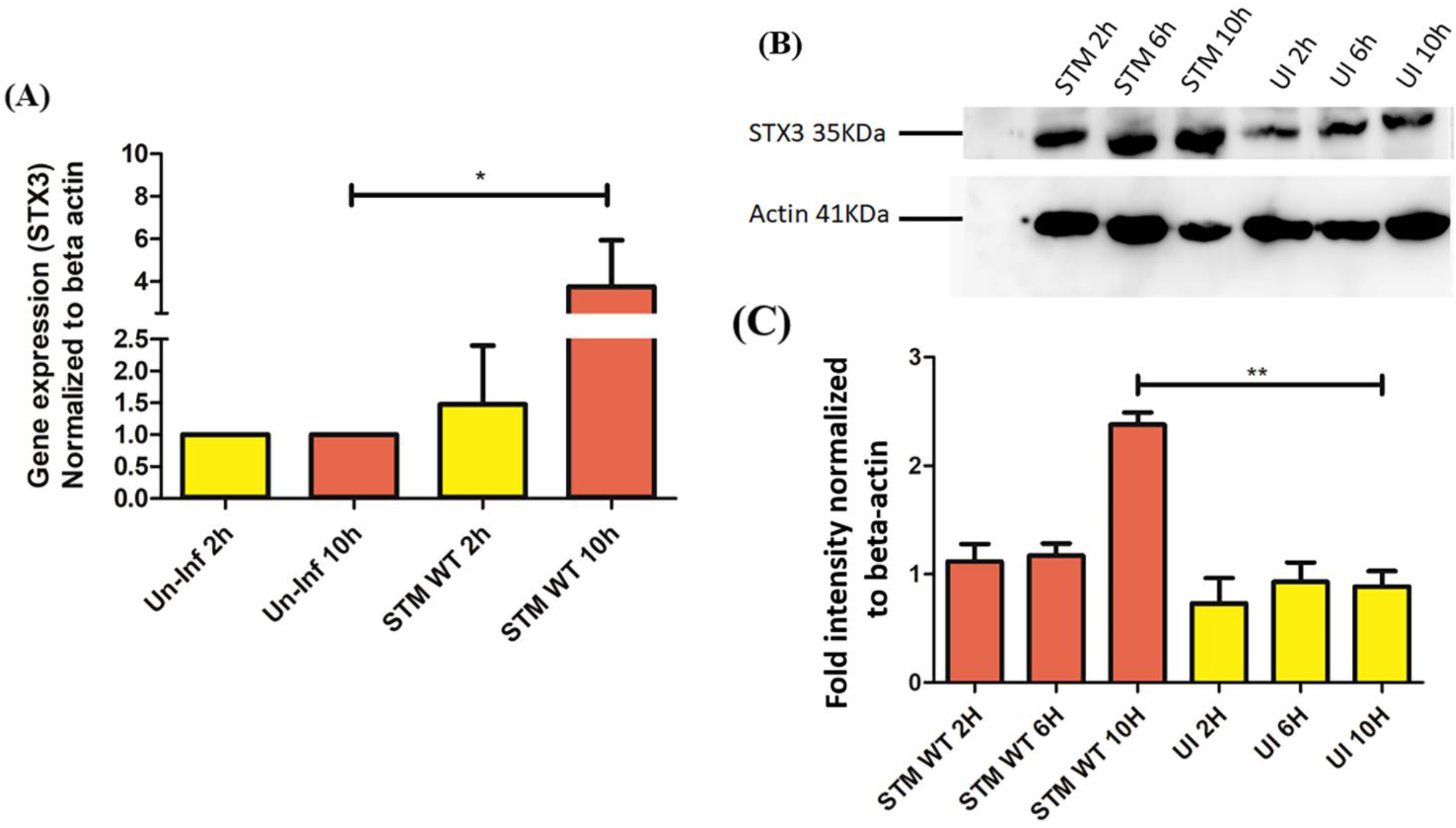
Upon STM infection both transcript and protein level of STX3 is up-regulated. (A) qRT-PCR at 2h and 10h time point, (B) Western blotting from murine macrophages (RAW264.7) infected with STM and uninfected at 2h, 6h and 10 timepoint, and (C) Graph plot for densitometric analysis done using ImageJ Platform. (N=3, n=3) (p<0.05-*, p<0.01-**, p<0.001-***).

### Live cell analysis reveals that STM WT maintains an association with STX3 until the late hours of infection

We wanted to decipher the interaction of STX3 with STM inside SCV at different time-point of infections. Therefore, we performed live cell imaging with GFP-STX3 transfected cells to monitor interactions and time-dependent changes upon infection with STM WT. We observed that the association of SCV with GFP-STX3 at early (2-4 hours) and intermediate (5-7 hours) time points post-infection was significantly higher as compared to late time-point (12-16 hours) **(Fig. 5 A)**. We have also taken control as non-pathogenic bacteria such as *E. coli* DH5α to decipher if this phenotype is *Salmonella* induced/infection specific. We observed that the association of STX3 with STM WT is significantly higher as compared to *E. coli* **(Fig. 5A-B)**. Our live cell imaging data suggest that STX3 maintains constant association with/localization to SCV till late time points of infection **(Fig. S5)**.

**Figure 5:**
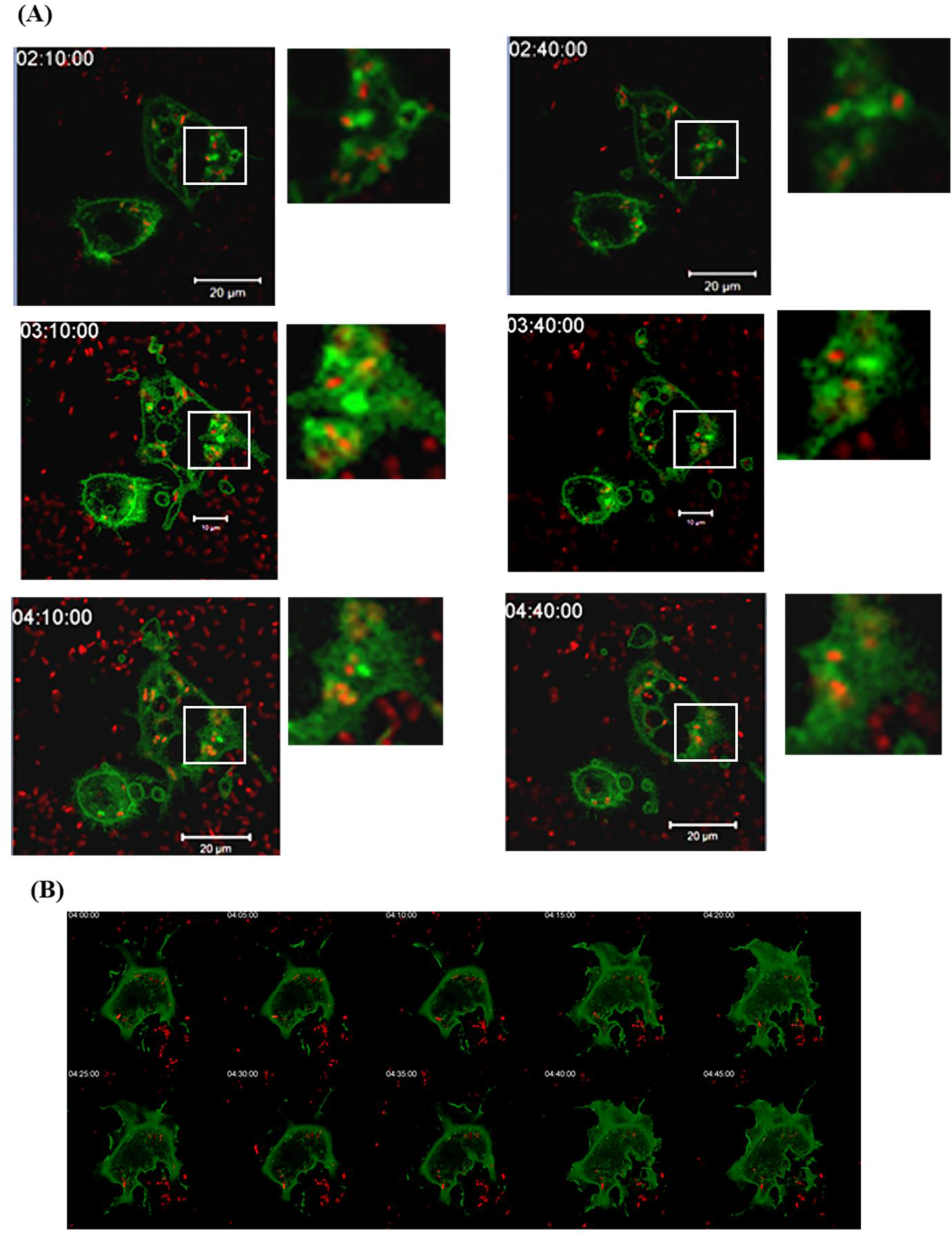
Live cell imaging. (A) Representative snapshot of live cell imaging of RAW264.7 murine macrophages expressing EGFP-STX3 and infected with mCherry expressing STM WT at an MOI-30, insets show constant contact between SCV and STX3; B) Representative snapshot of live cell imaging of RAW264.7 murine macrophages expressing EGFP-STX3 and infected with mCherry expressing *E. coli* at an MOI-30, we can see there is no contact between the bacteria and STX3 as a negative control.

### SPI-2 dependent regulation of STX3 acquisition to SCV in both *in-vitro* host cell infection and *in-vivo* mouse model

Next, we sought to decipher whether the colocalization that we observed during our live cell imaging experiments between SCV and STX3 are SPI-1 or SPI-2 dependent. We have therefore used apparatus knockout of SPI-1 (STM Δ*invC*) and SPI-2 (STM Δ*ssaV*) and compared the interactions with the STM WT infection in a time-dependent manner in RAW264.7 murine macrophages expressing EGFP-STX3. We observed that the association of SCV with STX3 are SPI-2 mediated, and the knockout of the SPI-2 apparatus leads to the abrogation of the colocalization between SCV and STX3 **(Fig. 6A-C)**. However, there is no significant difference between SPI-1 apparatus knockout and STM WT as per the colocalization coefficient quantification suggested using Zen 2.3**(Fig. 6A, C)**. STX3 has been reported to be present in the intestinal epithelial cells and has a significant role in maintaining the polarity of the cells[30]. There could be a possible role of STX3 in facilitating bacterial infection in the mice model of STM since enterocytes are one of the prime cell targets for STM. Therefore, to further validate our findings in an *in-vivo* mouse model of STM infection, we gavage the C57BL/6 mice with STM WT, STM Δ*invC* and STM Δ*ssaV*, isolated the ileum, performed sectioning and stained with anti-STX3 antibody and we found similar observation in *in-vivo* as well. The STM WT maintains association with STX3 as seen in the cross-sectional immune staining of distal ileum at 6h post gavaging from the mice gavage with STM WT and STM Δ*invC*. However, we could not observe any association of STX3 in the intestinal ileum at 6h post gavaging in the mice gavaged with STX3 Δ*ssaV*. Together these data indicate that the acquisition of STX3 to SCV and *in-vivo* ileum is dependent on SPI-2 system of STM **(Fig. 6E-F)**.

**Figure 6:**
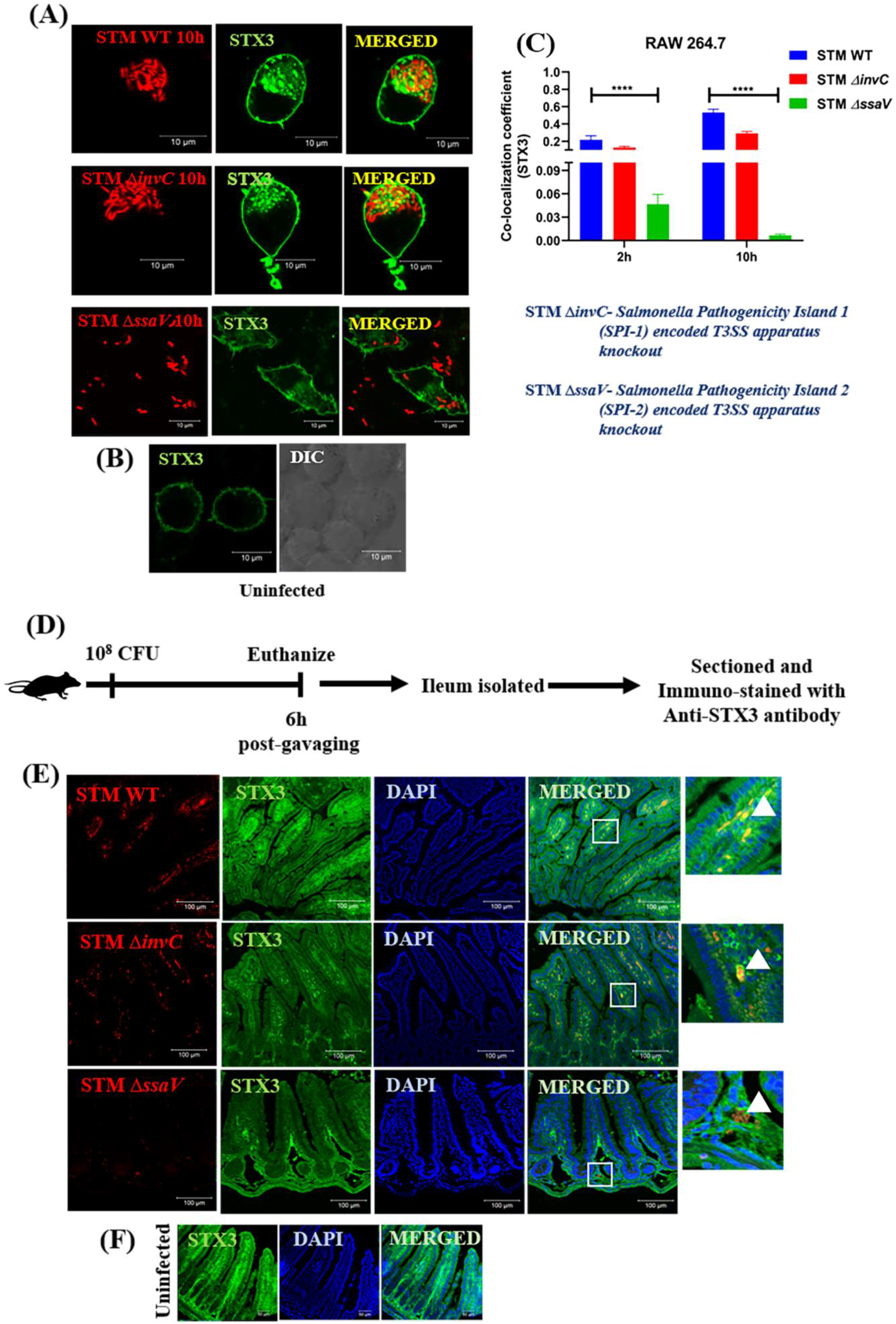
SPI-2 dependent regulation of STM with STX3 acquisition to SCV *in-vitro* and *in-vivo* mouse model. A) Representative immunofluorescence confocal images with RAW264.7 murine macrophages cells expressing EGFP STX3 infected at 10h post-infection with STM WT, STM Δ*invC* and STM Δ*ssaV;* B) Representative immunofluorescence confocal images with RAW264.7 murine macrophages cells expressing EGFP STX3 uninfected cells; D) Quantitation of colocalization coefficient at 2h and 10h post-infection in RAW264.7 macrophages in (A) and bacteria were individually marked using the ZEN 2.3 platform, and the data is a representative plot of three independent biological replicates. (N=4, n=3) (Student’s unpaired t-test was used p<0.05-*, p<0.01-**, p<0.001-***, p< 0.0001-****). D) Schematic of the *in-vivo* experiment protocol; E) Representative immunofluorescence confocal images with cross-sectioning of ileum staining with anti-STX3 antibody of mice gavaged with mCherry expressing bacterial cells (STM WT, STM Δ*invC* and STM Δ*ssaV)* after 6h post gavaging; F) Representative immunofluorescence confocal images with cross-sectioning of ileum staining with anti-STX3 antibody of mice gavaged with PBS (as uninfected control) after 6h post gavaging.

## Discussion

In eukaryotes, membrane fusion and fission are fundamental biological processes that assist in organelle biogenesis, secretion of molecules and uptake of various nutrients. It also facilitates important immune functions, including ingestion and destruction of invading pathogens[31]. Cargo transport between the organelles inside the cell is carried out by vesicles, which are assigned to deliver a cargo (proteins/lipids) and other bio-molecules from one compartment to another. During the intracellular transport, the membrane fusion events are regulated by specialized proteins called SNARE (soluble N-ethylmaleimide-sensitive factor attachment receptor) proteins. Through general phagocytosis or bacterial induced phagocytosis, intracellular pathogens gain access to host cells. Most of these successful pathogens, such as *Mycobacterium, Salmonella, Chlamydia* or *Legionella*, are capable of residing in a favorable compartment for survival, multiplication and establishment of pathogenesis. Bacteria need to maintain stable host niches, and therefore, these pathogens modify or alter the vesicle fusion events involving SNAREs to block the degradative fusion events, and acquire vesicles for various nutrients and host membranes. Our study observed that upon knocking down STX3, the number of bacteria per host cell is significantly reduced at 10 hours compared to untransfected or scrambled control. These suggest a possible role of STX3 in the survival of *Salmonella* in SCVs. We further observed that in STX3 knockdown, the number of bacteria that reside in one bacterium per vacuole is altered, and there are more events of multiple bacteria in a vacuole. These results indicate that STX3 might be utilized by bacteria to acquire host membrane and therefore play a crucial in the division and establishment of replicative niches. We also observed that the levels of SNAREs are upregulated upon *Salmonella* infection, suggesting the possible role *Salmonella* infection in inducing the host expression of STX3. We observed using live-cell imaging that SCV acquires STX3 during infection, and thus might help in fusion of endocytic vesicles with SCVs to acquire membrane for facilitating the growth and division of SCV. We also found this acquisition abrogated when we infected with SPI-2 encoded T3SS apparatus mutant (STM Δ*ssaV*) but not with SPI-1 encoded T3SS (STM *ΔinvC*). Together, these results helped us to develop a working model **(Fig. 7)** that the effector molecule/s secreted through SPI-2 encoded T3SS is involved in inducing the host expression of STX3 followed by its recruitment to SCVs, which is essential to maintain *Salmonella* division with a principle single bacterium per vacuole.

**Figure 7:**
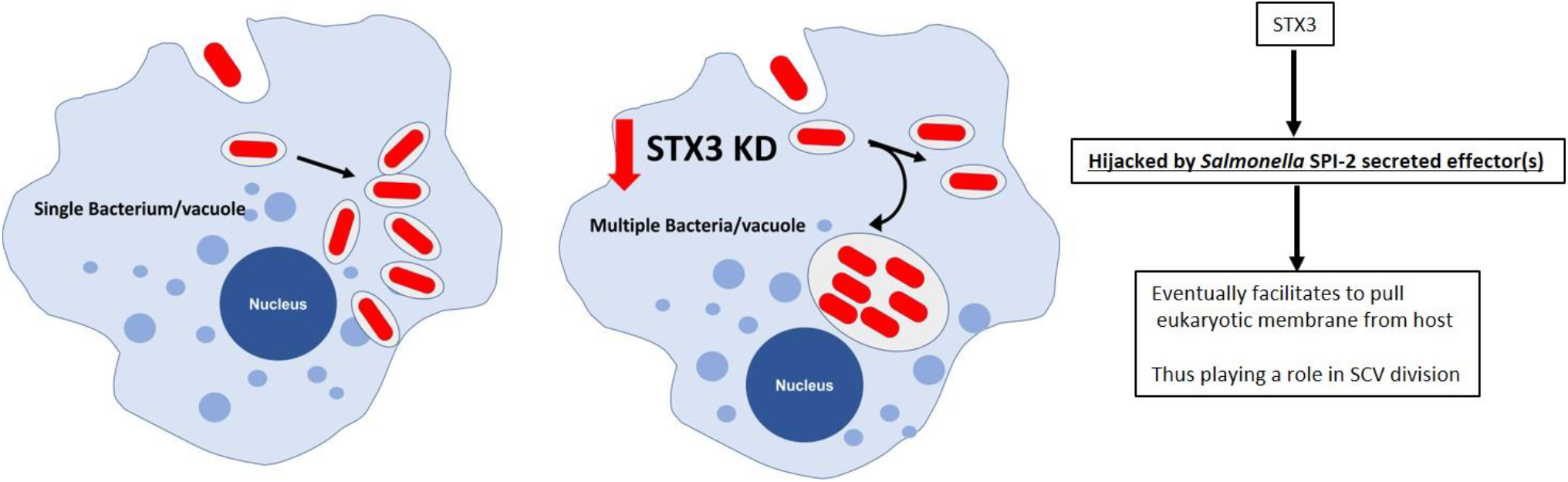
Schematic depicting the importance of STX3 in facilitating the replication of bacterium inside host cells.

## Materials and Methods

### Ethics statement

All animal experiments were reviewed and approved by the Institutional Animal Ethics Committee (IAEC) constituted as per article number 13 of the CPCSEA rules, laid down by the Government of India at the Indian Institute of Science, Bangalore INDIA (Acts, Rules and Amendments no:59 of 1960). IEAC Registration Number: 48/1999/CPCSEA; Project No: CAF/Ethics/854/2021.

### *In-vivo* experiments

All mice used (C57BL/6) were bred and housed at the Central Animal Facility, Indian Institute of Science, Bangalore, India. 4-6 weeks old, C57BL/6 mice were infected by oral gavaging of 10^8^ CFU of STM WT, STM *ΔinvC or* STM *ΔssaV*, 6 hours post-infection, mice were sacrificed, and Peyer’s patches from ileum section of intestine were isolated under aseptic conditions and stored in PFA until further histological processing. The samples were sectioned and immune-stained. The specimens were observed post staining with anti-STX3 antibody ((MAB2258, Merck), goat anti-mouse antibody) using Zeiss LSM 880 confocal laser scanning microscope, and the images were analyzed using the ZEN software.

### shRNA selection

To ensure maximum knockdown, we have targeted the 3’UTR of mRNA transcript of syntaxin 3, syntaxin 4, and syntaxin 12. We have used the Clustal-Omega web tool to align and identify a complementary target sequence to 3’UTR. shRNA is in TRC2-pLKO-puro vector (SHC201 Sigma-Aldrich) background with puromycin as a mammalian selection marker. Since human syntaxins have approx. 98% homology with mouse syntaxins, the same shRNA was used for RAW264.7.

### Bacterial strains and growth conditions

*Salmonella enterica* serovars Typhimurium wild-type (STM WT) strain ATCC 14028 or ATCC 14028 and the isogenic STM Δ*invC* (SPI-1 T3SS deficient) and STM Δ*ssaV* (SPI-2 T3SS deficient) mutants were used constitutively expressing either green fluorescent protein (GFP) or red fluorescent protein (mCherry) through pFPV25.1 were used in all experiments and in the study. *E. coli* DH5α is harboring the pLKO.2 plasmid encoding shRNA. All the bacterial strains were cultured in Luria broth (LB) with constant shaking (175rpm) at 37°C. Media was supplemented with ampicillin (50μg/ml) or kanamycin (50μg/ml) wherever required.

### Cell culture protocol

The cells RAW264.7 murine macrophages were cultured in DMEM - Dulbecco’s Modified Eagle Medium (Sigma) supplemented with 10% FBS at 37°C in a humidified incubator with 5% CO2. Prior to each experiment, the cells were seeded onto the required plate either with a coverslip (for confocal fluorescence microscopy) or without (for intracellular survival assay) at a confluency of 50-60%.

### Transfection protocol

PEI-mediated transfection was carried out wherein the plasmid DNA harboring shRNA targeted syntaxin 3 (STX3), syntaxin 4 (STX4), syntaxin 12 (STX12) and scramble shRNA (SCR) in a concentration of 300 ng/well was incubated along with PEI in a 1:2 ratio for 20 mins in serum-free DMEM. Following this, we added the concoction to mammalian cell systems such as RAW264.7 after 6-8 hours; the media was changed with DMEM + 10% FBS. After 48h, the cells were used for different experimental setups.

### Gentamicin protection assay

The transfected cells were then infected with *Salmonella* Typhimurium (STM) tagged with or without red fluorescent protein (mCherry) at MOI of 25 for the confocal experiment. Upon infecting the RAW264.7 cell line with STM-mCherry/STM, the plate was centrifuged at 700-900 rpm for 5 mins to facilitate the adhesion and then incubated for 20mins at 37°C and 5% CO_2_. Post-incubation, the bacteria-containing media were removed, wells were twice washed with PBS, and fresh media containing 100µg/mL gentamicin was added and incubated for 1 hour at 37°C and 5% CO_2_. Following this, the media was removed, washed with PBS twice, and 25µg/mL of gentamicin-containing media was added and incubated at 37°C and 5% CO2 for different time points. The time points selected for confocal microscopy were 2 hours, 6 hours and 10 hours post-infection.

### Confocal Microscopy

After appropriate hours of incubation post-infection with STM-WT-GFP, the cells on coverslips were washed thrice with PBS and fixed with 3.5 % paraformaldehyde for 10-15mins. Then cells were washed twice with PBS and incubated with a specific antibody (α-LAMP1) in a blocking buffer containing 2 % BSA and 0.01% saponin for 3 hours at room temperature (RT) or overnight at 4°C. Following this, the cells were washed twice with PBS and incubated with an appropriate secondary antibody tagged with fluorochrome for 1 hour at RT. The coverslips were then mounted onto a clean glass slide with mounting media and antifade agent; after the mounting media dried, it was sealed with clear nail polish and imaged under a confocal microscope.

### RNA isolation and quantitative RT PCR

RNA isolation was performed from transfected cells / after appropriate hours of infection with STM WT at MOI of 10 using TRIzol (Takara) reagent according to manufacturers’ protocol. Quantification of the RNA was done in NanoDrop (Thermo-Fischer scientific). To check for RNA quality, the isolated RNA was also run on 2% agarose gel, and 3μg of RNA has subjected to DNase 1 treatment at 37°C. The reaction was then stopped with the addition of EDTA, heating the sample at 65°C for 10mins. The cDNA was synthesized by incubating the isolated DNA-free RNA with oligo (dT)18, and 5X RT buffer, RT enzyme, dNTPs, and DEPC treated water at 42°C for 1 hour. Quantitative real-time PCR was done using SYBR green RT-PCR kit in BioRad qRT-PCR system. All the reaction was set up in a 384 well plate with three replicates for each sample. The gene expression levels of interest were measured using specific RT primers. Gene expression levels were normalized to beta-actin as an internal control.

### Immunoblotting

After appropriate hours of infection with STM WT at MOI of 10, the media was removed, and the cells were washed twice with PBS. Cells were then harvested using a sterile scraper and centrifuged at 1500 rpm for 10 mins, 4°C. Cell lysis was done by RIPA buffer for 30mins on ice, followed by estimation of total protein using the Bradford protein estimation method. Polyacrylamide Gel Electrophoresis (PAGE) was done by loading 35μg of protein from whole cell lysate, then transferring it onto 0.45μm PVDF membrane (GE Healthcare). The membrane was blocked using 5% skimmed milk (Hi-Media) in TTBS for 1h at RT and was then probed with specific primary and secondary HRP conjugated antibodies. The membrane was developed using ECL (Bio-rad), and images were captured using ChemiDoc GE healthcare. All densitometric analysis was performed using the Image J Platform.

### Live cell imaging

Cells were seeded onto a glass bottom live cell imaging dish (Eppendorf) at a confluency of less than 50%; 12 hours later, the cells were transfected using FuGENE HD or lipofectamine 3000 (as per manufacturer’s protocol) and pEGFP-C1 plasmid encoding Rat Syntaxin 3 (EGFP-STX3) was used. 48h post-transfection, cells were infected with STM WT or other mutants expressing mCherry at an MOI of 30 and incubated at 37°C and 5% CO2 for 30mins; cells were then washed twice with PBS, and fresh DMEM medium containing 25μg/mL gentamicin was added, and imaging was performed in LSM 710 Zeiss microscope at 37°C and 5% CO2 and 63X oil immersion objective, till the end of time-points.

### Statistical Analysis

Statistical analyses were performed with GraphPad Prism software. The Student’s t-test was performed as indicated. The results are expressed as mean ± SD or mean ± SEM. Group sizes, experiment number, and p values for each experiment are described in figure legends.

## Acknowledgment

Divisional and Departmental Confocal imaging Facility, shRNA resource centre, Departmental Real-Time PCR Facility, and Central Animal Facility at IISc are duly acknowledged. Mr. Punith, Mrs. Saima and Ms. Navya are acknowledged for their help in image acquisition.

## Availability of data and materials

All data generated and analysed during this study, including the supplementary information files, have been incorporated in this article. The data is available from the corresponding author on request.

