## supplementary data for "Syntaxin 3-SPI 2 dependent cross-talk facilitates the division of *Salmonella* containing vacuole (SCV)"

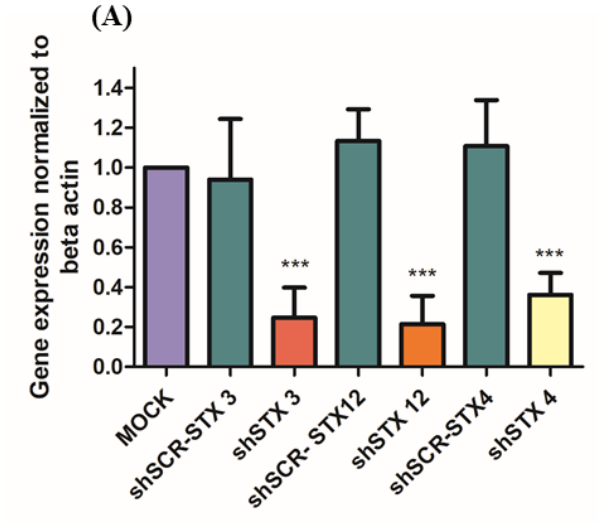


**Figure S1: Knockdown of host STX3, 4 and 12 does not affect phagocytosis of *S.* Typhimurium in murine macrophages RAW 264.7.** Representative images of knockdown macrophages infected with *Salmonella* Typhimurium at 2 hours post infection. (A) Mock treated, (B) shSCR (scrambled), (C) shSTX3, (D) shSTX4 , (E) shSTX12 and (F) Quantification of number of bacteria/host cell. (N=3, n= 50 microscopic field) ($p<0.05$- *, $p<0.01$- **, $p<0.001$- ***).

S2

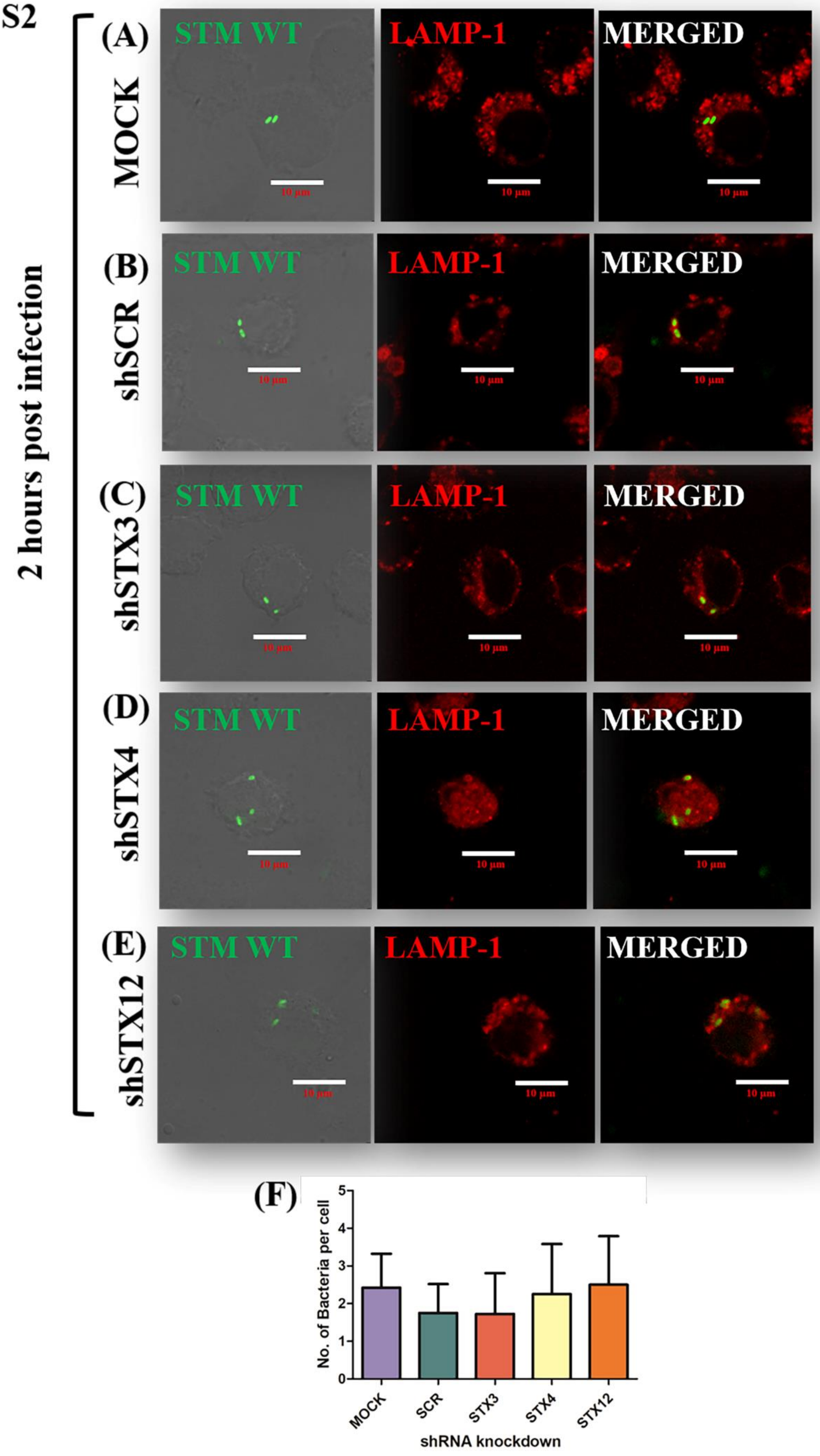
2 hours post infection
(A)
MOCK
(B)
shSCR
(C)
shSTX3
(D)
shSTX4
(E)
shSTX12
STM WT
LAMP-1
MERGED
10 µm
(F)
No. of Bacteria per cell
0
1
2
3
4
5
MOCK
SCR
STX3
STX4
STX12
shRNA knockdown

**Figure S2: Knockdown of host STX3, 4 and 12 leads to reduced no. of bacteria/ host cell at 2 hours post infection with STM WT (GFP) in murine macrophages RAW 264.7.** Representative images of knockdown macrophages infected with *Salmonella* Typhimurium at 6 hours post infection. (A) Mock treated, (B) shSCR (scrambled), (C) shSTX3, (D) shSTX4, (E) shSTX12 and (F) Quantification of number of bacteria/host cell. (N=3, n= 50 microscopic field) ($p<0.05$- *, $p<0.01$- **, $p<0.001$- ***).

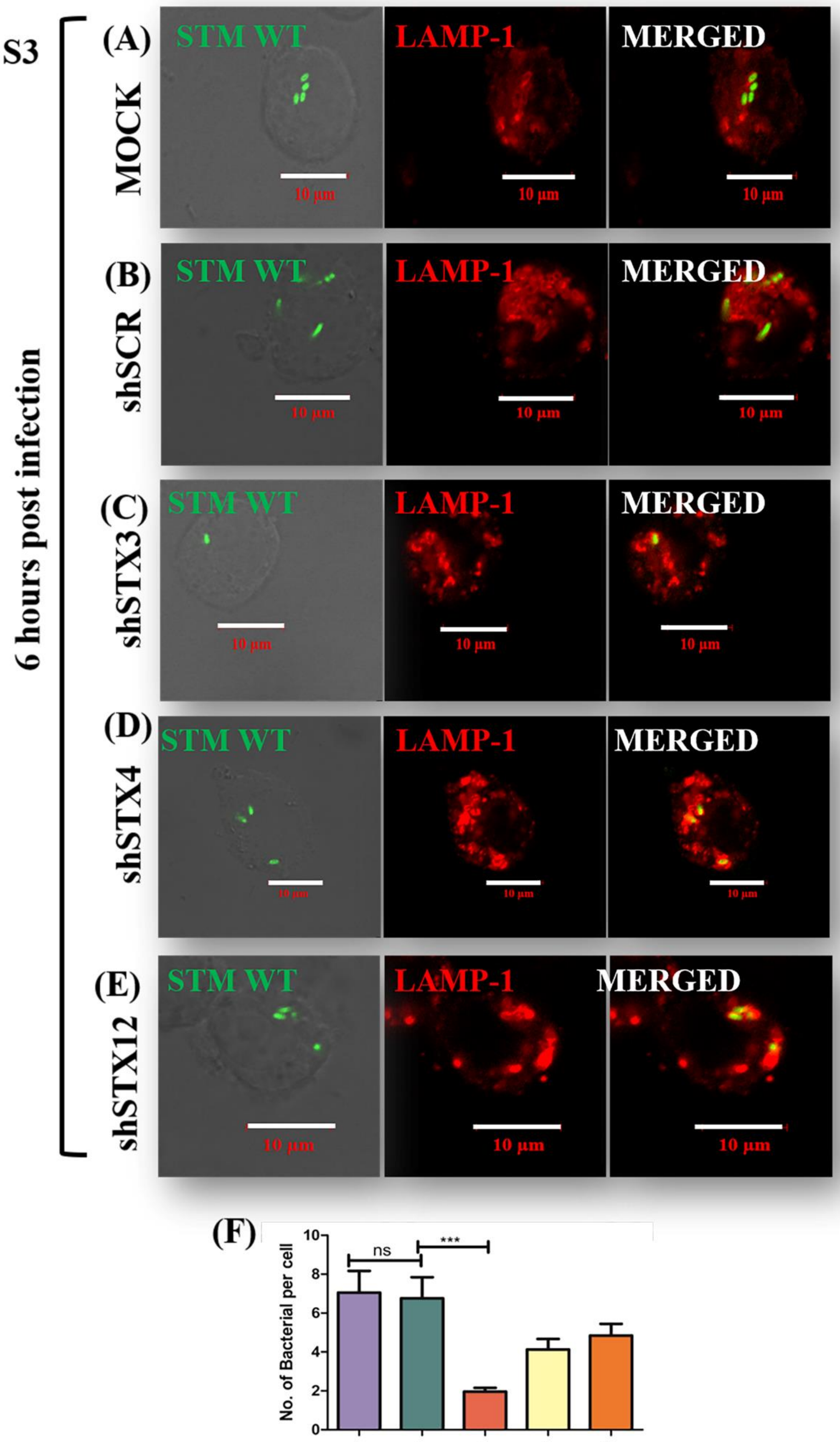
S3
6 hours post infection
(A)
MOCK
(B)
shSCR
(C)
shSTX3
(D)
shSTX4
(E)
shSTX12
STM WT
LAMP-1
MERGED
10 µm
(F)
No. of Bacterial per cell
ns
***
MOCK
sh SCR
sh STX3
sh STX4
sh STX12
shRNA knockdown

**Figure S3: Knockdown of host STX3, 4 and 12 leads to reduced no. of bacteria/ host cell at 6 hours post infection with STM WT (GFP) in murine macrophages RAW 264.7.** Representative images of knockdown macrophages infected with *Salmonella* Typhimurium at 6 hours post infection. (A) Mock treated, (B) shSCR (scrambled), (C) shSTX3, (D) shSTX4 , (E) shSTX12 and (F) Quantification of number of bacteria/host cell. (N=3, n= 50 microscopic field) ($p<0.05$- *, $p<0.01$- **, $p<0.001$- ***).

S4

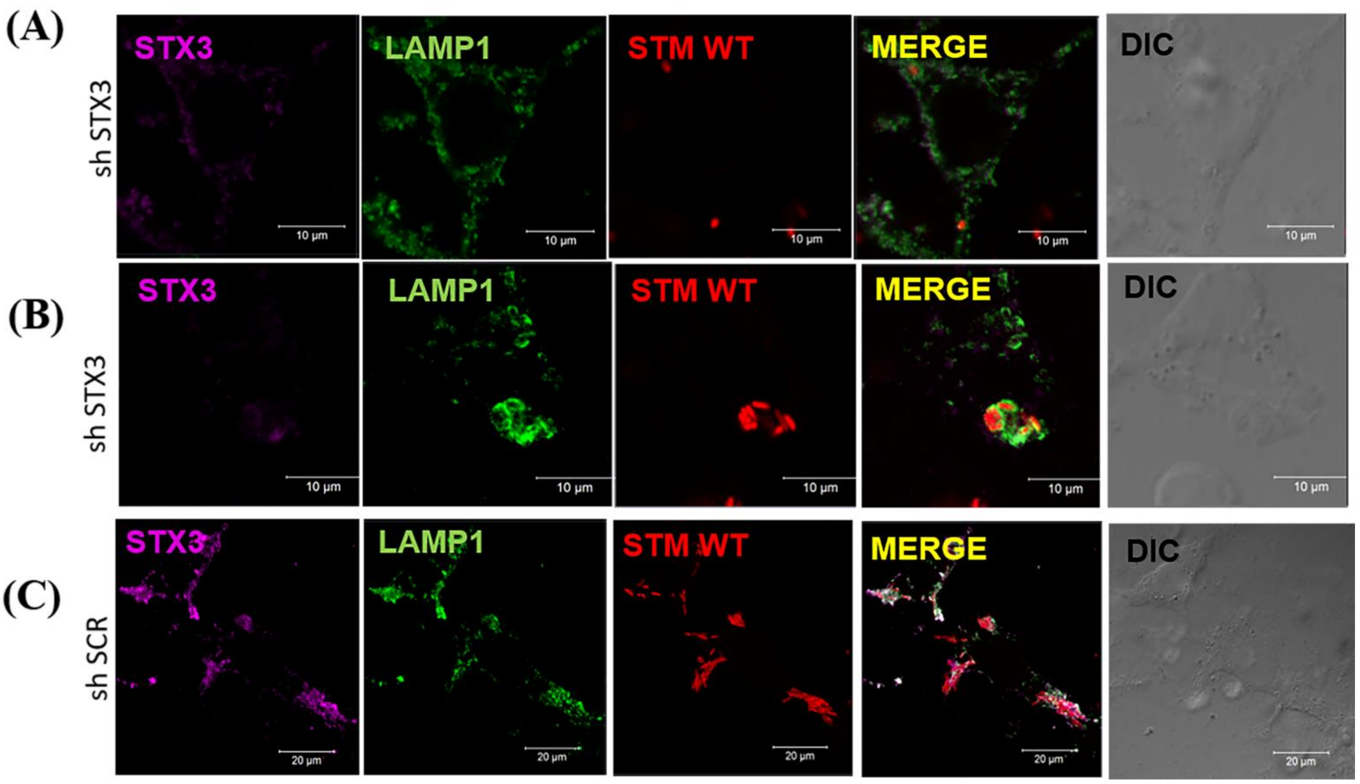


**Figure S4: Reconfirmation of STX3 knockdown phenotype of multiple bacteria in a single vacuole.** (A) Representative confocal microscopy images of RAW264.7 cells transfected with shSTX3 and infected with STM WT and further stained with anti-STX3 antibody less number of bacteria/ host cells; (B) Representative confocal microscopy images of RAW264.7 cells transfected with shSTX3 and infected with STM WT and further stained with anti-STX3 antibody multiple bacteria in a single vacuole; (A) Representative confocal microscopy images of RAW264.7 cells transfected with shSCR and infected with STM WT and further stained with anti-STX3 antibody

(A)

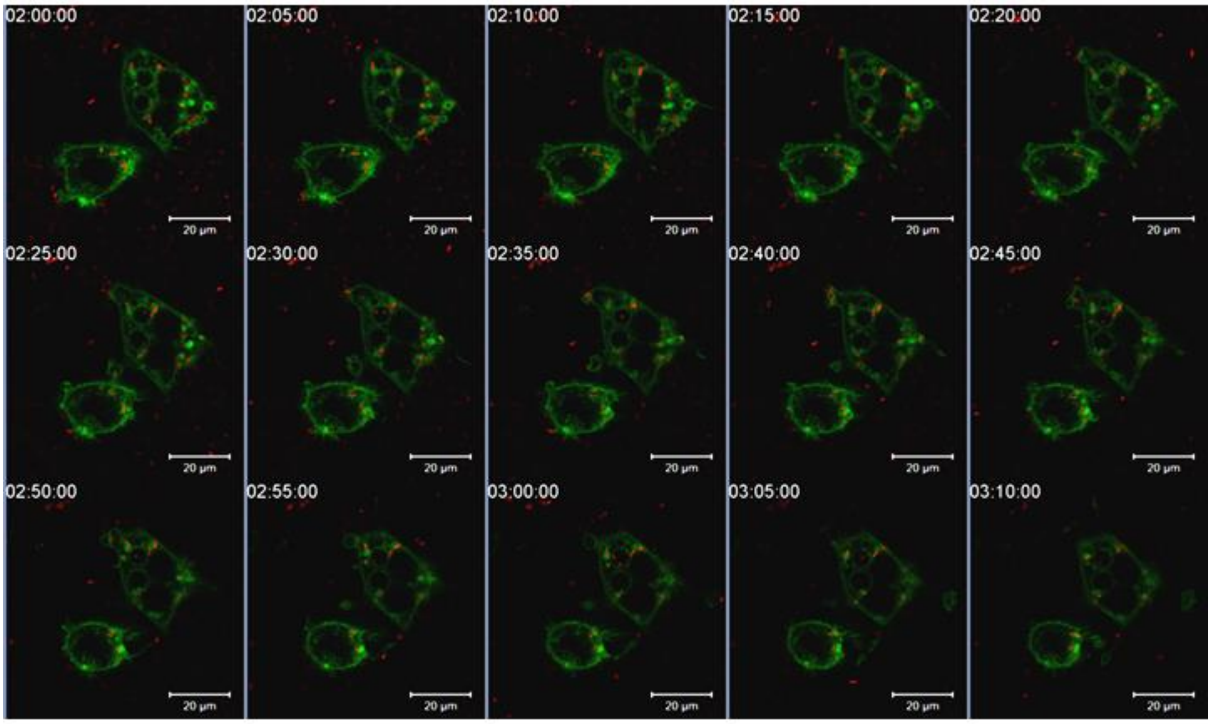
02:00:00
02:05:00
02:10:00
02:15:00
02:20:00
02:25:00
02:30:00
02:35:00
02:40:00
02:45:00
02:50:00
02:55:00
03:00:00
03:05:00
03:10:00
20 μm

(B)

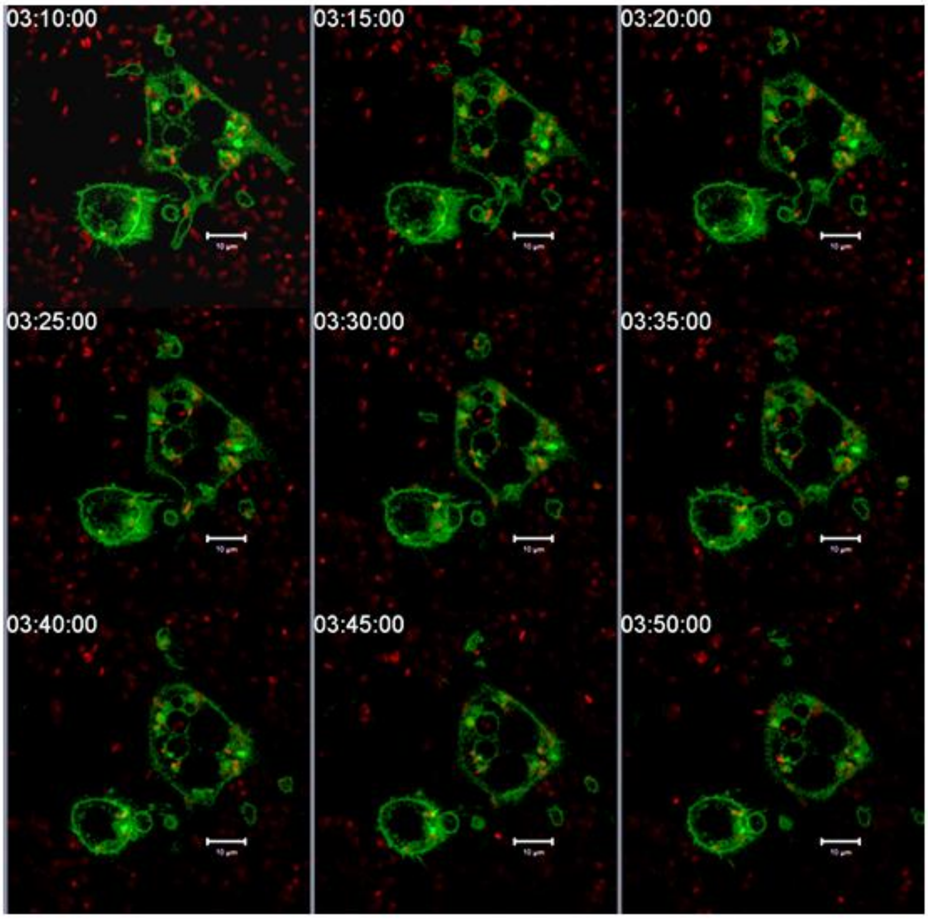
03:10:00
03:15:00
03:20:00
03:25:00
03:30:00
03:35:00
03:40:00
03:45:00
03:50:00

(C)

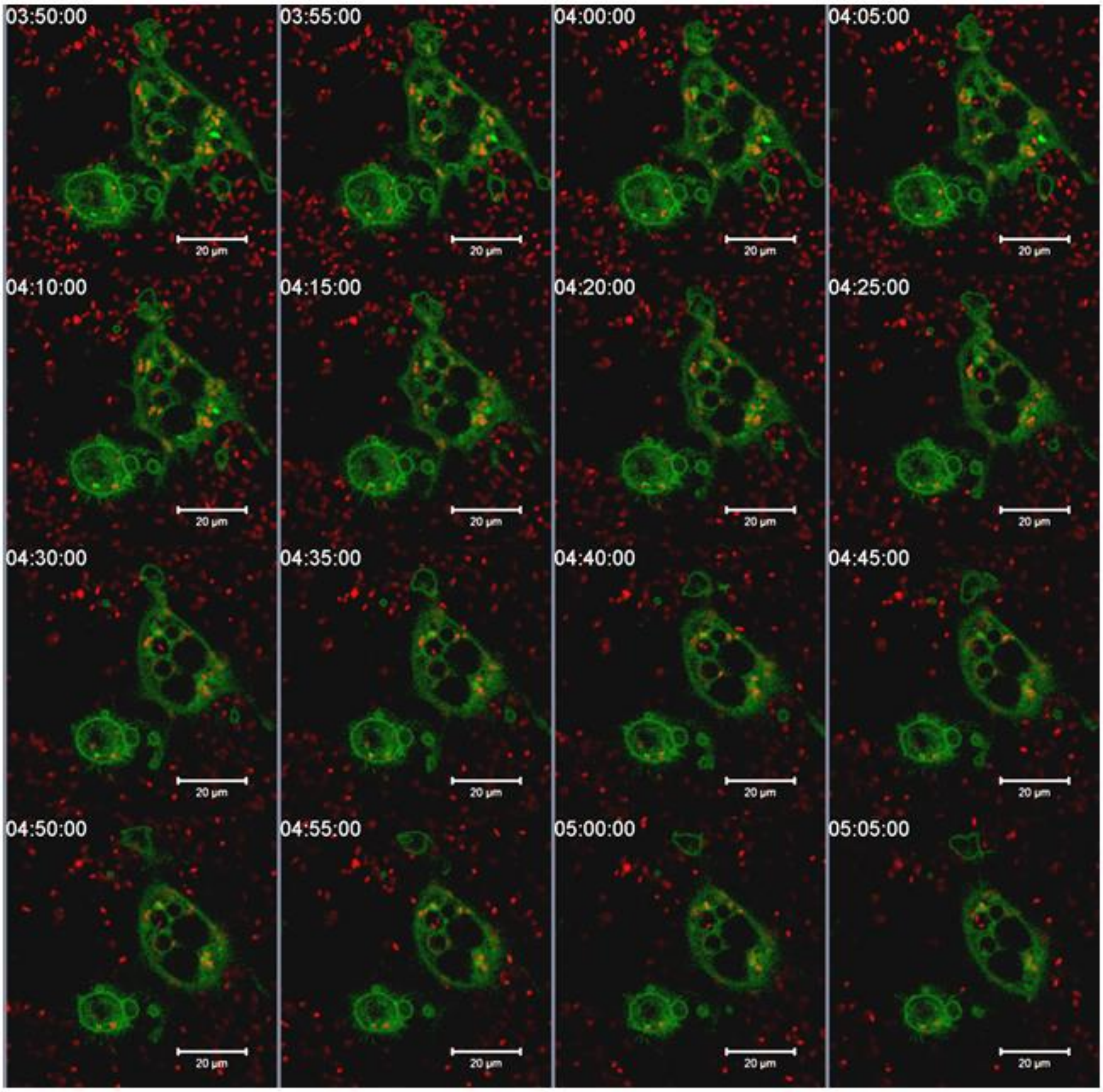
03:50:00
03:55:00
04:00:00
04:05:00
04:10:00
04:15:00
04:20:00
04:25:00
04:30:00
04:35:00
04:40:00
04:45:00
04:50:00
04:55:00
05:00:00
05:05:00
20 µm

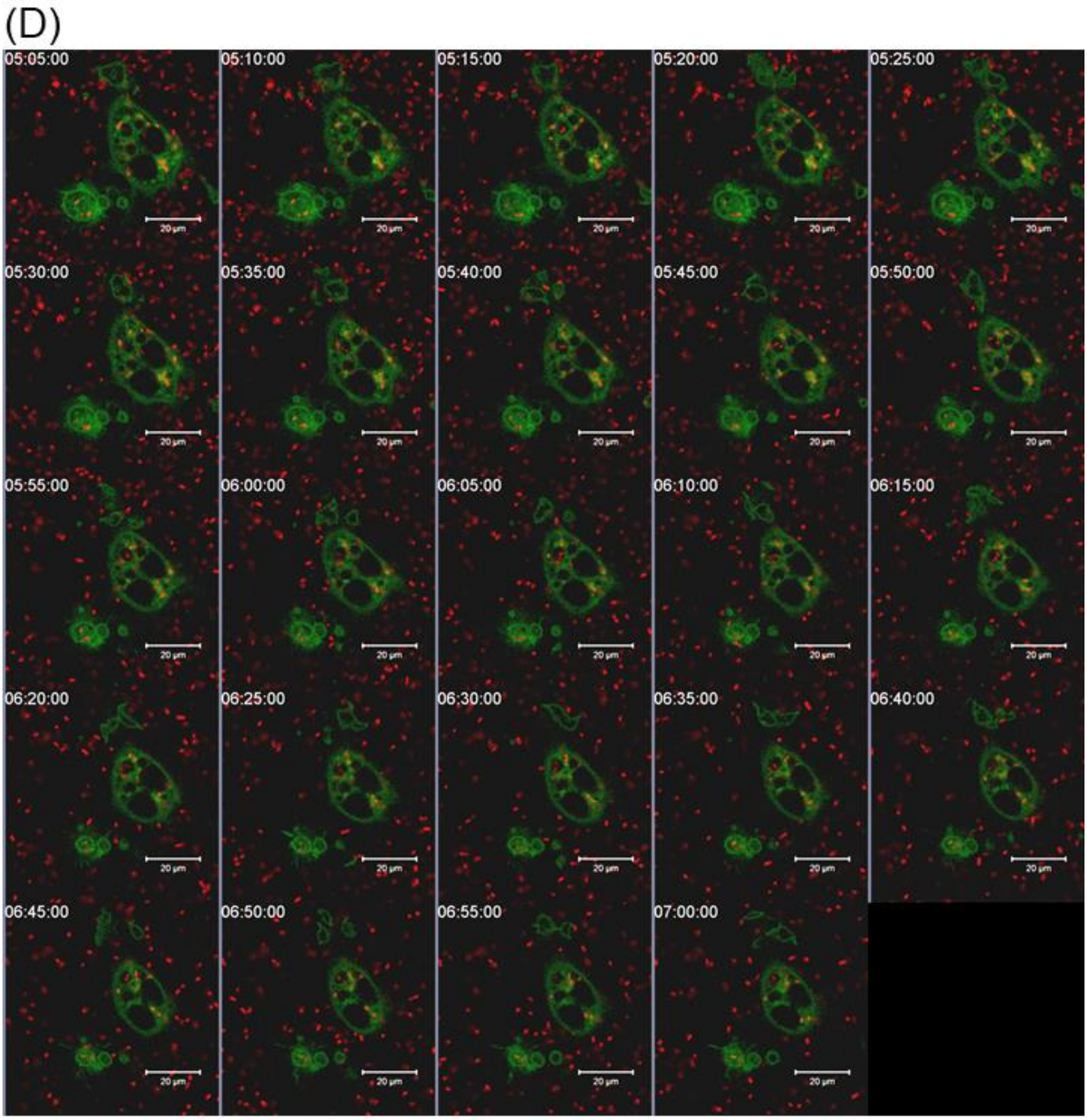


**Figure S5: Time series snap shots of the live cell imaging**

A-D Representative snapshot of live cell imaging of RAW264.7 murine macrophages expressing EGFP-STX3 and infected with mCherry expressing STM WT at an MOI-30, showing constant association between SCV and STX3;
